# An ecological model of masting reproduction matches empirical dynamics

**DOI:** 10.64898/2026.09.02.748907

**Authors:** Mahdi Salehzadeh, John M. Stockie, Ailene MacPherson

## Abstract

Masting, characterized by highly variable, synchronized, and intermittent seed or fruit production, represents a common reproductive strategy among perennial plants and has profound ecological consequences. Resource provisioning and pollen limitation have long been viewed as central physiological mechanisms underlying this strategy, recent empirical evidence also highlights the role of weather cues in initiating and synchronizing reproductive effort. Drawing on mechanisms that drive periodicity in disease dynamics, this study proposes an alternative proximate mechanism for masting. We develop and analyze a stage-structured population growth model in which developmental delays create population-level cycles, and demographic stochasticity adds individual-level variation; together, yielding masting-like patterns. We compare the behaviour of this novel model with that of the widely used resource budget model and empirically observed patterns of masting in perennial plants. To quantify and compare model outputs and empirical observations, we employ three continuous metrics of masting that capture volatility, synchrony, and periodicity. Our study provides an alternative proximate mechanism for masting. Comparison of this novel mechanism and the established resource-budget model to empirical time-series reveals that both represent realistic yet distinct forms of masting reproduction. Together, these models provide a foundation for further exploration of the conditions under which this reproductive strategy can evolve. Beyond masting, our results highlight the general importance of life-history timing and demographic stochasticity in shaping population ecology.

## 1 Introduction

Reproductive effort in perennial plants is often highly variable between years and synchronized among individuals, a phenomenon known as masting [Kelly, 1994, Pesendorfer et al., 2021]. Masting plays a crucial role in a wide variety of ecological processes, triggering pronounced boom-and-bust cycles that impact food webs, influence plant growth, affect animal migrations, and even have implications for human disease dynamics [Clark et al., 2019, Lauder et al., 2019, Zuckerberg et al., 2020, Bregnard et al., 2021, Jourńe et al., 2023]. A comprehensive description of masting requires examining both the proximate (e.g., physiological and demographic) and ultimate (e.g., eco-evolutionary) mechanisms underlying this reproductive strategy [Kelly, 1994, Pesendorfer et al., 2021, Bogdziewicz et al., 2024a]. Understanding these mechanisms is significant for ecologists, foresters, land managers, and conservation biologists, particularly given growing evidence that climate change is altering the frequency and intensity of mast events [Foest et al., 2024, Bogdziewicz et al., 2024b, Pearse et al., 2021]. In the current study, we focus on mechanistic explanations for masting and investigate under what conditions individual-level processes, such as resource allocation, life-history structure, and demographic stochasticity, scale up to produce variability, synchrony, and periodicity in reproductive effort, and how well these mechanisms align with long-term empirical data.

Resource allocation has been established as a possible ultimate mechanism for masting. In the resource-budget framework [Isagi et al., 1997, Satake and Iwasa, 2000, 2002b, Venner et al., 2016, Pearse et al., 2016], individual plants accumulate resources annually and reproduce only when their resources exceed a threshold, resulting in interannual variation in reproductive output. When coupled with population-wide pollen limitation, this model can also synchronize reproductive effort among individuals [Satake and Iwasa, 2000, 2002a]. Intermittent, synchronous reproduction may be further reinforced by ecological or life-history traits, such as disturbance-mediated cues (e.g., canopy gaps or fire) [Tachiki and Iwasa, 2008, Beck et al., 2024], multi-year juvenile stages [Tachiki and Iwasa, 2010], and broader trait-based patterns associated with masting [Pearse et al., 2020, Jourńe et al., 2023]. Extensions of the resource budget model incorporating pollen limitation have been used to examine when mechanisms such as pollination efficiency or predator satiation can favour the evolution of masting [Tachiki and Iwasa, 2010, 2012, 2013]. This approach has provided important insights into the ecological and evolutionary basis of masting. Nevertheless, it remains uncertain whether this model is the sole mechanistic explanation for this reproductive strategy or whether alternative processes may also contribute to masting dynamics.

Evaluating and comparing these proximate mechanisms requires a common framework for quantifying masting which captures both the magnitude and the temporal structure of reproduction in a multidimensional way. Reproductive effort often varies on a continuum, rather than switching discretely between “mast” and “non-mast” years [Kelly, 1994, Crone et al., 2011, Bogdziewicz et al., 2024a]. Hence, we adopt the continuous, spectral and correlation-based metrics summarized by Qiu et al. [2023], which capture three non-mutually exclusive features of reproductive dynamics extracted from time series of reproductive output (A). They include: (1) “volatility” in individual reproductive success as determined by the variability in individual seed set over time; (2) “synchrony”, which refers to the tendency of individuals to produce large seed crops in the same years; and (3) “periodicity”, representing the interval between years of high seed production at a population level. These metrics reflect both an individual reproductive behaviour and an emergent population-level phenomenon. Periodicity in life-history events is not unique to masting; for instance, infectious diseases often exhibit recurrent outbreaks or irregular epidemics. Mechanisms such as seasonality, quarantine, age or stage structure, and the presence of multiple pathogen strains can explain these sustained cycles in infection prevalence [Hethcote, 2000, Nuño et al., 2005, Goņcalves et al., 2011, MacPherson and Otto, 2018]. We draw on this stage-structured epidemiological perspective (e.g., Susceptible vs. Infected vs. Recovered) to motivate a mechanism for population-level periodic reproduction, one key descriptor of masting. Classic compartmental models implicitly assume the transition between epidemiological stages (e.g., transition from infected to recovered) is captured by an exponential distribution of waiting times (e.g., the duration of infection is exponentially distributed). While this simplifying assumption is often sufficient to capture key epidemiological features (e.g., the basic reproductive ratio (R_0_)), it is not biologically realistic as the exponential distribution has a mode of zero, implying that most individuals transition immediately (e.g., instantaneous recovery from infection). More realistic transition timings or “delays” can be modeled by introducing sequential “sub-stages”, producing Erlang-distributed waiting times; this principle is formalized in the linear chain trick and its extensions [Lloyd, 2001, Hurtado and Kirosingh, 2019]. Beyond epidemiology, reproduction in perennial plants provides a natural example of such life history delays. Before producing a large seed crop, individuals must sequentially accumulate resources, initiate floral primordia, develop buds, and mature seeds—each step requiring a characteristic duration. Modeling adults as progressing through a chain of reproductive sub-stages, therefore, offers a biologically grounded representation of plant physiology.

Masting is defined not only by population-level periodicity but also by significant inter-annual variation in individual reproductive effort (volatility) and synchrony among individuals. These individual-level characteristics arise when stochastic reproduction of individuals within a population are linked through shared population level mechanisms such as crowding and competition. This coupling aligns reproductive pulses across individuals while maintaining substantial year-to-year variability. Building on established epidemiological and ecological models, here we introduce a stage-structured model that incorporates realistic life-history timing and demographic stochasticity, and explicitly tracks both individual- and population-level seed production as a potential alternative proximate mechanism for masting. Masting-like patterns may also result from environmental forcing, such as seasonality. Similarly seasonality can affect epidemic outbreaks, yet many baseline epidemic models assume no seasonal forcing. Consistent with this approach, the present analysis focuses on the non-seasonal case to demonstrate that life-history timing and demographic stochasticity alone are sufficient to generate masting-like patterns.

In the current study, we develop a stage-structured framework as an alternative proximate mechanism for masting. We begin, however, with a review of the resource-budget model with pollen limitation to facilitate the comparison of this established mechanism and our novel mechanism. While motivated by epidemiological models, the proposed stage-structured model for masting is based on incorporating biologically realistic life-histories in which individuals vary relatively little (compared to the classic exponential model) in the duration of each life stage and the incorporation of stochasticity in individual reproductive events. Using a deterministic modeling framework, we first consider under what conditions the stage-structured model is expected to exhibit cyclic population dynamics indicative of periodic reproductive effort. To incorporate individual-level aspects of masting, we then introduce a stochastic model which incorporates effects of demographic stochasticity arising from individual reproductive timing and seed set. We conclude by comparing both the resource-budget and the stage-structured mechanisms for masting to observed patterns of reproductive effort in published time series of reproductive output obtained predominantly from tree species. Taken together, this work demonstrates that common features of individual life histories (stage-structure and aging) and population growth (density-regulation and stochasticity) can, in combination, produce masting-like reproductive effort. This novel mechanism for mating broadens theoretical avenues for investigating ultimate, evolutionary explanations for this reproductive strategy.

## 2 Review: Resource-Budget Model

First introduced by Isagi et al. [1997], the resource–budget model has become a dominant proximate explanation for masting, supported by empirical studies [Venner et al., 2016, Schermer et al., 2019, Ito and Kudo, 2024]. The resource-budget model characterizes how an individual plant allocates resources to reproduce. Each mature plant accrues an annual net production from photosynthesis, denoted by *P_s_*. Reproduction is triggered once the available reserve surpasses a threshold, *L_T_*, for flowering and fruiting. Specifically, let *R_i_*(*t*) be the resource level of individual plant *i* at the beginning of year *t*. If *R_i_*(*t*) + *P_s_ < L_T_*, the plant does not reproduce that year and carries its resources forward. If *R_i_*(*t*) + *P_s_* ≥ *L_T_*, the plant allocates the surplus to flowering and additional resources to fruiting. In the basic model, the “flowering cost” equals the excess above threshold, *C_f_* (*t*) = *R_i_*(*t*) + *P_s_* − *L_T_*, and the “fruiting cost”, which is used as a measure of reproductive effort, is proportional to the flowering cost, *C_a_*(*t*) = *R_c_ C_f_* (*t*), where *R_c_* is the proportionality constant. Thus, the post-reproduction resource is reduced by both *C_f_* (*t*) and *C_a_*(*t*). The one-year update for the individual-level resource-budget model is:

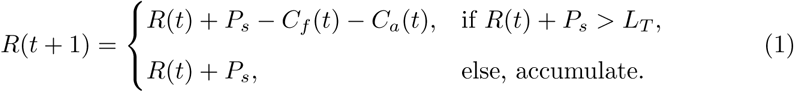

Model parameters and the ranges explored are summarized in Table 1; throughout, we adopt the baseline values and exploration ranges from Isagi et al. [1997]. As *R_c_* increases beyond one, the model exhibits amplification of small differences in resource levels, causing individual reproductive trajectories to diverge over time (Fig. 1a). In addition, we quantify this transition using the volatility metric, which captures interannual variability in reproductive effort (Fig. 1b).

**Fig. 1:**
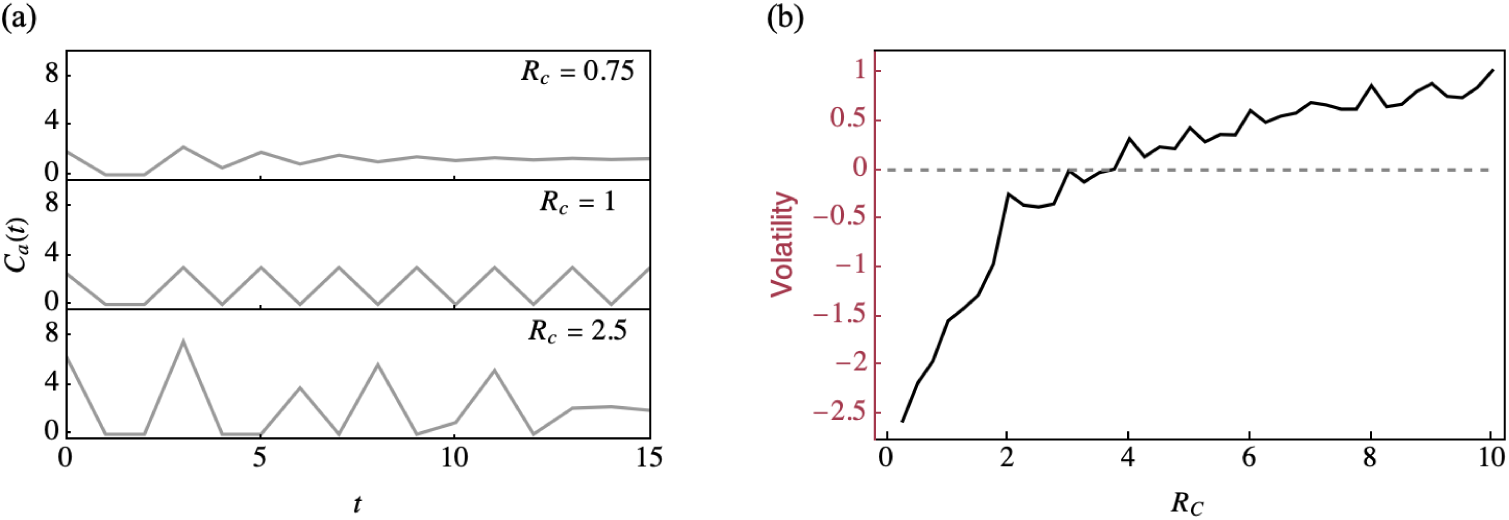
Resource-budget model at individual-level. (a) Time-series of annual fruiting cost *C_a_*(*t*), used in the resource-budget model as the measure of reproductive output, for various ratios of fruiting to flowering costs, *R_c_*. (b) Volatility (log(V*_x_*)) in annual fruiting cost *C_a_*(*t*) as a function of *R_c_* while other parameters are held constant. *P_s_*, *L_T_* and *C_f_* (0) were 3, 6 and 2.5 respectively.

**Table 1:**
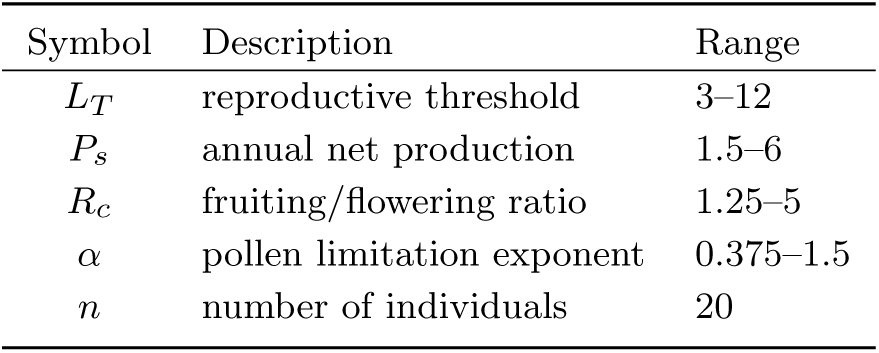
Summary of resource–budget model parameter ranges we tested.

| Symbol | Description | Range |
| --- | --- | --- |
| $L_T$ | reproductive threshold | 3–12 |
| $P_s$ | annual net production | 1.5–6 |
| $R_c$ | fruiting/flowering ratio | 1.25–5 |
| $\alpha$ | pollen limitation exponent | 0.375–1.5 |
| $n$ | number of individuals | 20 |

Synchronous reproduction among individuals arises in this model through the introduction of pollen limitation in the form of a density-dependent fruiting term [Isagi et al., 1997]. By assuming that the fruiting cost for individual *i* scales with the population’s flowering proportion via an exponent *α*, then reproductive synchrony in a population with *n* individuals can be written as

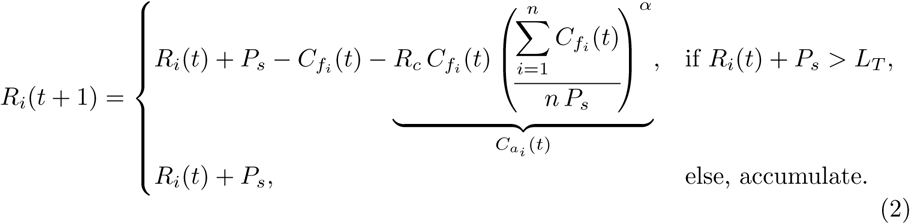

Here, pollination (and thus fruiting efficiency) increases with the population’s flowering proportion, and *α* controls how strongly this proportion modifies fruiting cost, a form of density-dependence, through ecological mechanisms such as pollinator behaviour, and pollen limitation. Different values of *α* determine whether synchrony in flowering emerges gradually or abruptly as the number of flowering individuals increases. Illustrative individual trajectories and synchrony as a function of *α*, are shown in Fig. 2a and 2b, respectively.

**Fig. 2:**
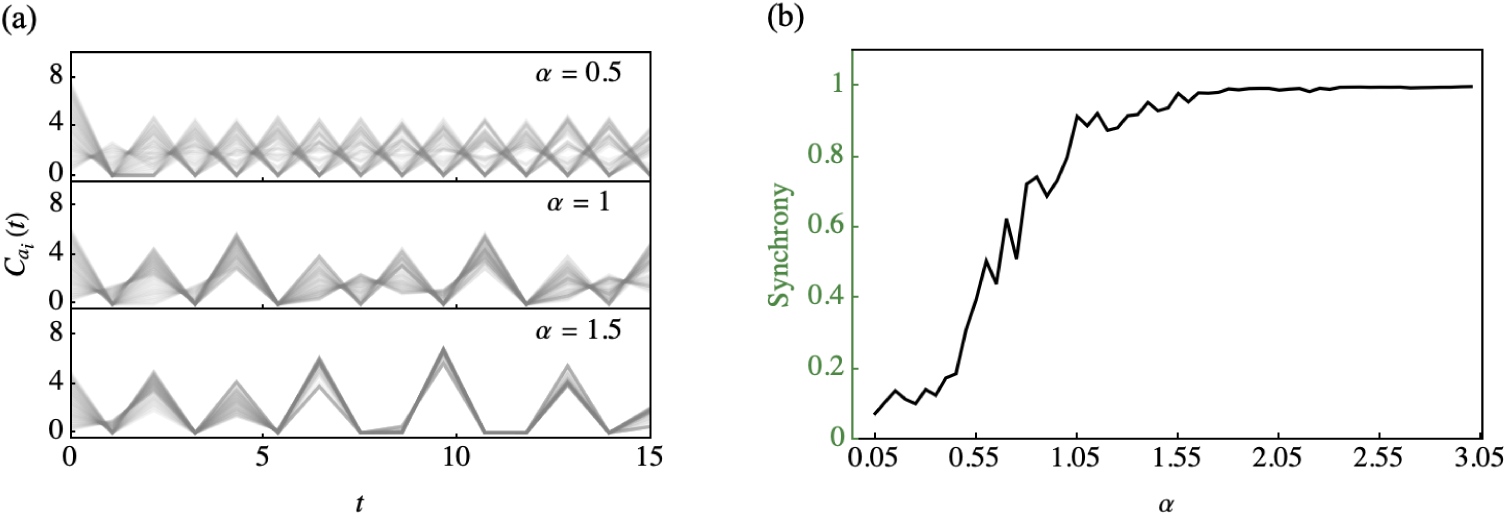
Resource–budget model at population-level. (a) Simulated time series of annual fruiting cost (*C_ai_* (*t*)) across different values of *α*. (b) Synchrony (S*_x_*) in annual fruiting cost *C_ai_* (*t*) as a function of *α*, with all other parameters held constant. *P_s_*, *L_T_*, *C_f_* (0) and *R_c_* were 3, 6, 1.5 and 2.5 respectively.

## 3 Stage-Structured Model

We develop a logistic population-growth model with juvenile and adult stages to study how life-history timing, together with density dependence and demographic stochasticity, can generate masting-like reproduction. In this approach, we first employ the deterministic model to demonstrate how stage structure can produce population-level cycles through the shifting concentrations of individuals between stages. Building on these results, we introduce an individual-based stochastic implementation and show that demographic noise generates persistent individual-to-individual heterogeneity that resembles masting-like variability and synchrony.

Following common practice in epidemiological modeling [Hethcote, 2000, Nuño et al., 2005, Gonçalves et al., 2011, MacPherson and Otto, 2018], we refine each lifehistory stage by introducing a sequence of sub-stages that allows us to manipulate development timing. Throughout, we refer to these as “sub-stages” for convenience, but they serve purely as a mathematical device to shape waiting-time distributions and do not necessarily correspond to distinct biological stages. Biologically, the juvenile stage represents immature individuals undergoing a delayed maturation process, whereas the adult stage represents reproductive individuals subject to turnover (mortality/senescence). Specifically, in a stage-structured model with a single juvenile and adult sub-stage and a constant rate of aging, the distribution of waiting times until adulthood (e.g., duration of the juvenile stage) will be exponentially distributed. By introducing *p* sub-stages within each life-history stage, we can model a more realistic Erlang distribution of waiting times. Specifically, juveniles progress through sub-stages *J*_1_*, . . ., J_p_* before reaching maturity, while adults transition through substages *A*_1_*, . . ., A_p_* before dying of old age (i.e., adult turnover). Hypothetically the number of stages could differ between juvenile and adult stages but we use the same number throughout for simplicity since previous epidemiological models have found that periodicity arises when the number of sub-stages is large in both sequential stages [Gonçalves et al., 2011, MacPherson and Otto, 2018]. Density dependence is incorporated as a general regulatory mechanism, such as competition, space, or resource limitation, which constrains population size and links individual reproductive dynamics through a common population density regulation.

A list of parameters and their illustrative values is provided in Table 2. This system structure is illustrated in Fig. 3a, and the following set of differential equations governs the population dynamics:

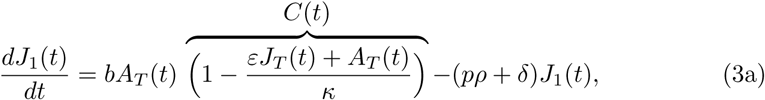

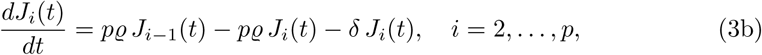

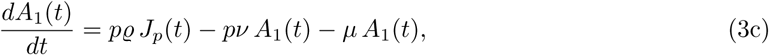

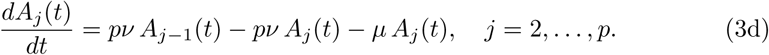

**Fig. 3:**
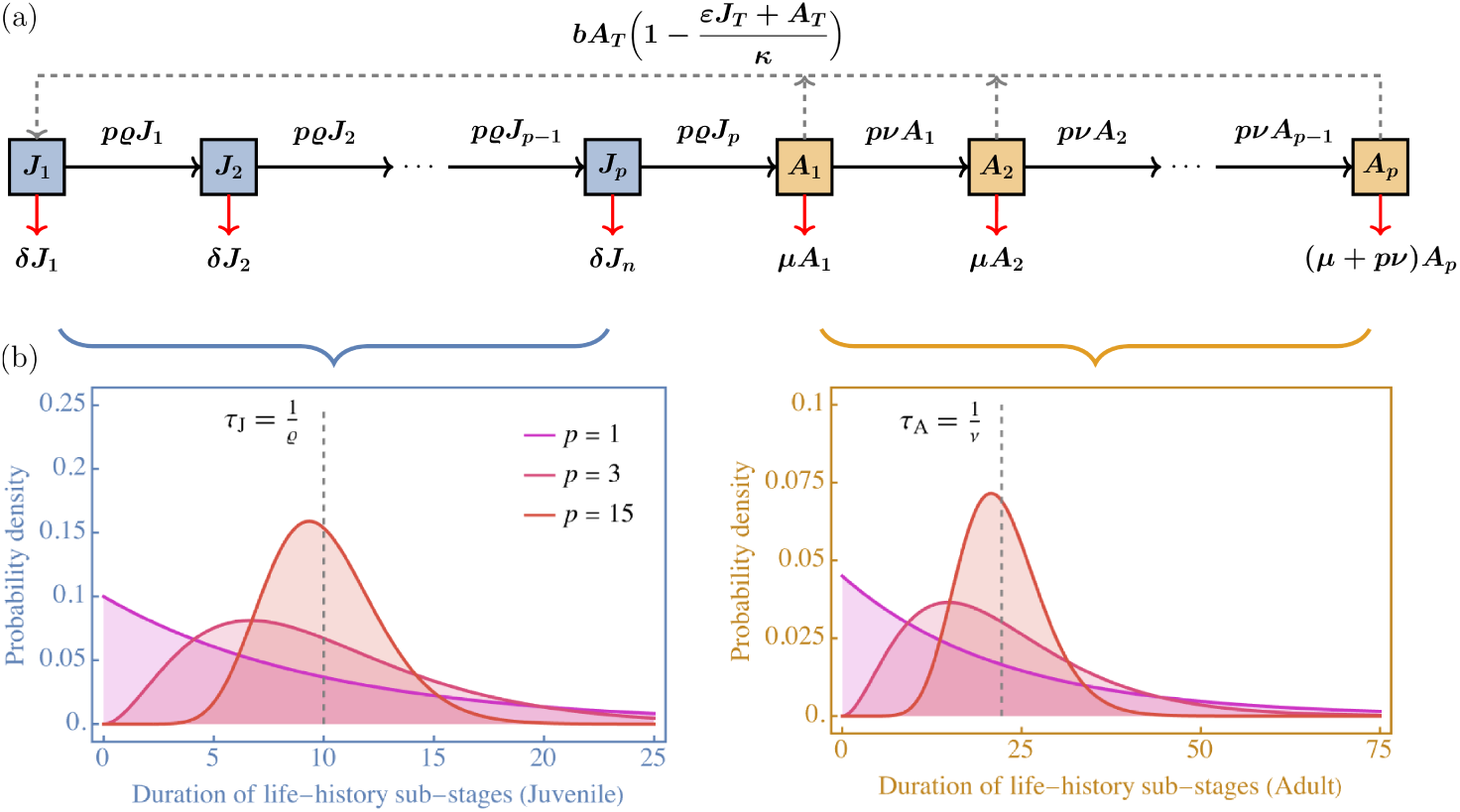
Stage-structured model and its implied waiting-time distributions. (a) Model schematic showing transitions among juvenile (*J_i_*) and adult (*A_j_*) substages with density-dependent reproduction. (b) Probability density distributions of sub-stage durations for juveniles and adults as the number of sub-stages, *p*, increases, demonstrating reduced temporal variability and convergence toward fixed mean durations.

**Table 2:** Summary of stage-structured model parameters and values explored.

| Symbol | Description | Values |
| --- | --- | --- |
| $p$ | number of sub-stages | 1–20 |
| $\varepsilon$ | juvenile competition weight | $10^{-4}$–1 |
| $b$ | per-capita reproduction rate | 0.2 |
| $\rho$ | juvenile maturation rate | 0.1 |
| $\nu$ | adult turnover rate | 0.045 |
| $\delta$ | juvenile mortality rate | 0.002 |
| $\mu$ | adult mortality rate | 0.001 |
| $\kappa$ | carrying capacity | 100 |

Here, adults reproduce at an intrinsic per-capita rate of *b*. Therefore, in the absence of density dependence, the total production of new juveniles is proportional to the total adult population, 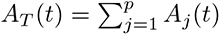. Let 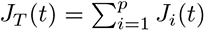 denote the total juvenile population size. Reproduction is reduced by density dependence scaling factor on reproduction, denoted as *C*(*t*), where *κ* is the carrying capacity and *ε* ∈ [0, 1] is the contribution of juveniles to crowding relative to adults. Thus, *C*(*t*) scales reproductive output according to the current total population size, representing general limitation due to competition for space and/or resources. Juveniles and adults are removed through mortality at rate *δ* and *µ* respectively. Juveniles mature through each of the *p* sub-stages at an exponential rate *pϱ*. As a result the expected time that an individual spends as a juvenile is 1*/ϱ* (in the absence of mortality) and is independent of the number of sub-stages *p*. Leaving the mean unchanged, the number of sub-stages, *p*, impacts only the variance in waiting times–low *p* results in high variability in the duration of the juvenile stage among individuals (see Fig. 3b). Similarly, adults advance through sub-stages at a rate *pν* such that the expected life span as an adult is 1*/ν* in the absence of background mortality.

### 3.1 Population-Level Deterministic Model

The deterministic stage-structured model exhibits a single, biologically-valid, nonzero “persistence” equilibrium. Because the persistence condition becomes analytically intractable for larger values of *p* we located the extinction boundary numerically by determining, for each *p*, the critical recruitment rate *b*_crit_(*p*) at which the dominant eigenvalue of the extinction equilibrium changes sign. Given that extinction does not occur (*b > b*_crit_(*p*)), we determined the stability of this persistence equilibrium numerically in the bifurcation diagram (Fig. 4). For *p >* 1 the leading eigenvalues are often complex, producing oscillatory transients about the persistence equilibrium. These cycles either decay toward the stable equilibrium (negative real part of the leading eigenvalue, points A & B in the bifurcation diagram, Fig. 4) or exhibit stable limit cycles (unstable persistence equilibrium with a positive real part of the leading eigenvalue, point C in the bifurcation diagram, Fig. 4). We confirmed the existence and stability of the limit cycles using numerical Floquet analysis (Appendix. B). The stability of the persistence equilibrium depends predominantly on the number of sub-stages (*p*) and the relative contribution of juveniles to density regulation (*ε*). Sustained limit cycles characteristic of population-level periodicity emerge under conditions of weak juvenile crowding (small *ε*) and low variance in development time (large *p*).

**Fig. 4:**
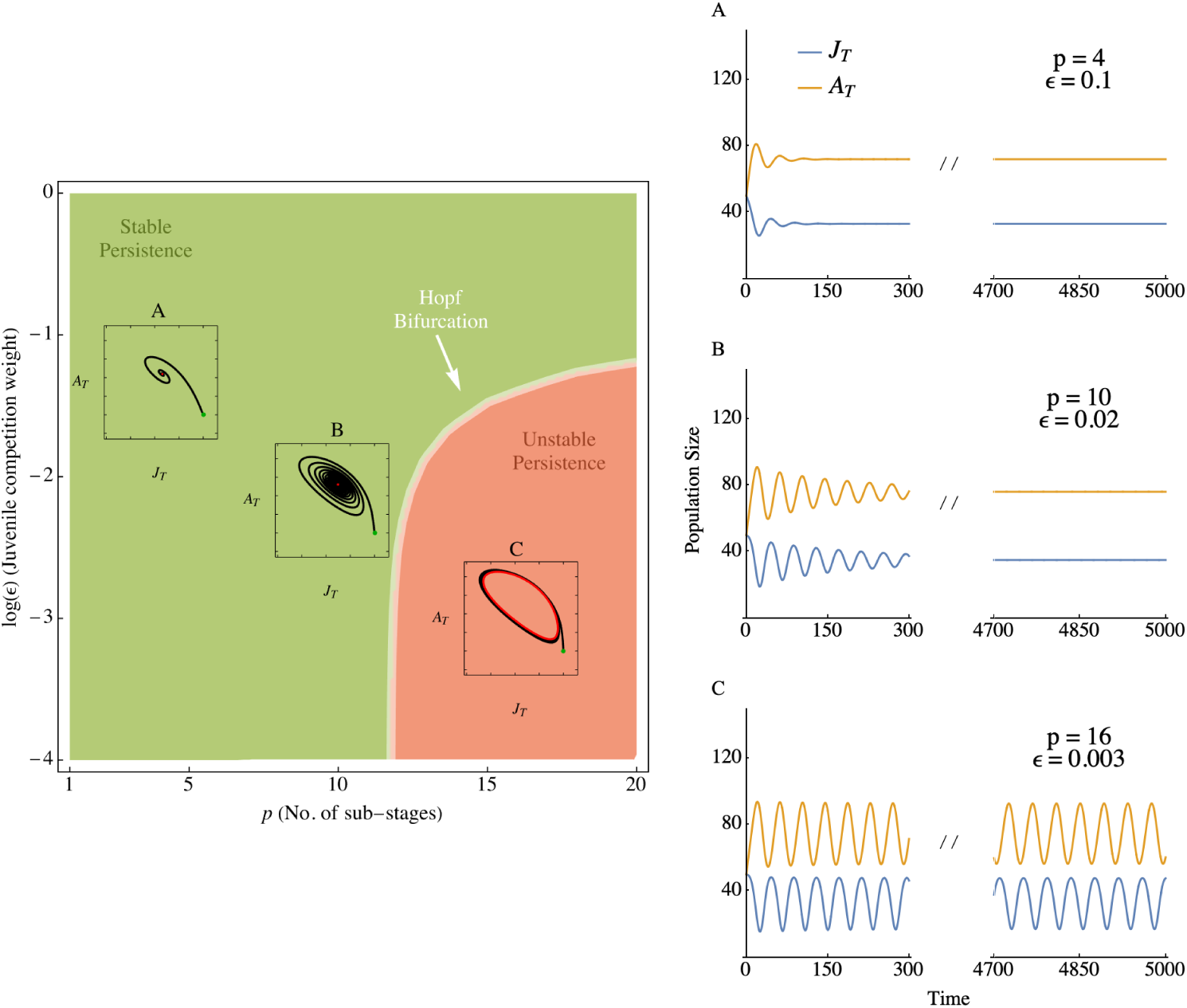
Emergence of limit cycles in the stage-structured model. Bifurcation diagram in (*p, ε*) parameter space with the Hopf bifurcation curve where Re[*λ*_max_] changes sign. Insets show representative phase portraits in the (*J_T_, A_T_*) plane for the parameter combinations (*p, ε*) corresponding to A:(4,0.1), B:(10,0.02), and C:(16,0.003). Green dots denote initial conditions. Red dots indicate stable persistence equilibria, and red closed curves indicate stable limit cycles. Panels A, B, and C show the corresponding deterministic dynamics of total juveniles, and adults. All other parameters are fixed as listed in Table 2, and with initial conditions *J_T_* (0) = 50 and *A_T_* (0) = 50.

As the distribution of development times narrows (large *p*), juvenile maturation and adult reproduction become increasingly clustered in time. When juvenile density dependence is weak (small *ε*), these clustered events destabilize the persistence equilibrium and give rise to stable limit cycles. This behavior is consistent with classical epidemiological results showing that clustered recovery (analogous to maturation in our model) often modeled via distributed delays, can induce recurrent outbreaks. Notably, population-level variability in the deterministic model does not originate from inter-annual variation among individuals. Empirical studies demonstrate that individual plants often display considerable year-to-year variability in reproductive output, and population-level fluctuations arise when this individual-level variability becomes synchronized across the population. In contrast, individuals in the deterministic model are identical and perfectly synchronized by design, so population-level volatility reflects coherent stage dynamics rather than the aggregation of heterogeneous individual behaviors. This distinction motivates the individual-based stochastic formulation introduced in the next subsection, where individual-level volatility arises endogenously and can scale up to population-level patterns through synchrony.

### 3.2 Individual-Based Stochastic Model

We simulate the stage-structured model as a continuous-time and discrete-space stochastic process using the Gillespie direct algorithm [Gillespie, 1976]. At any moment, several types of events can occur, including births, aging of juveniles and adults through the sub-stages, deaths in any sub-stage, and maturation of the oldest juveniles into the adult chain. Each event type has a rate that depends on the current abundance of juveniles and adults (Table 1 of Supplementary Material; Algorithm 1).

Beyond stochasticity in event timing, the model incorporates stochasticity in seed set at each birth event. Conditional on a birth event occurring at time *t*, the number of seeds produced by the reproducing adult, Δ*J*_1_, is given by the following sigmoid function:

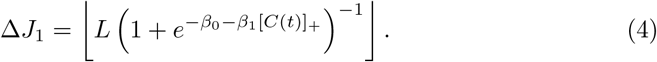

Here [*C*(*t*)]_+_ represents the nonnegative value of the density-dependent crowding term (i.e., [*a*]_+_ = max{0*, a*}), *L* is the maximum seed size, and *β*_0_ = −4.595, *β*_1_ = 9.190 are fixed so that seed size spans nearly the full admissible range, [0.01*L,* 0.99*L*], over the biologically relevant levels of crowding (Fig. 1 of Supplementary Material). Biologically, this reflects the assumption that seed set increases with access to available resources and declines under stronger competition, while remaining bounded above by *L*.

Fig. 5 shows a representative realisation for the parameter set corresponding to case C in Fig. 4. At the population level, the model exhibits recurrent fluctuations in total juvenile and adult population size. At the individual level, it generates heterogeneous but temporally clustered seed production time series, producing masting-like patterns of volatility, synchrony, and periodicity.

**Fig. 5:**
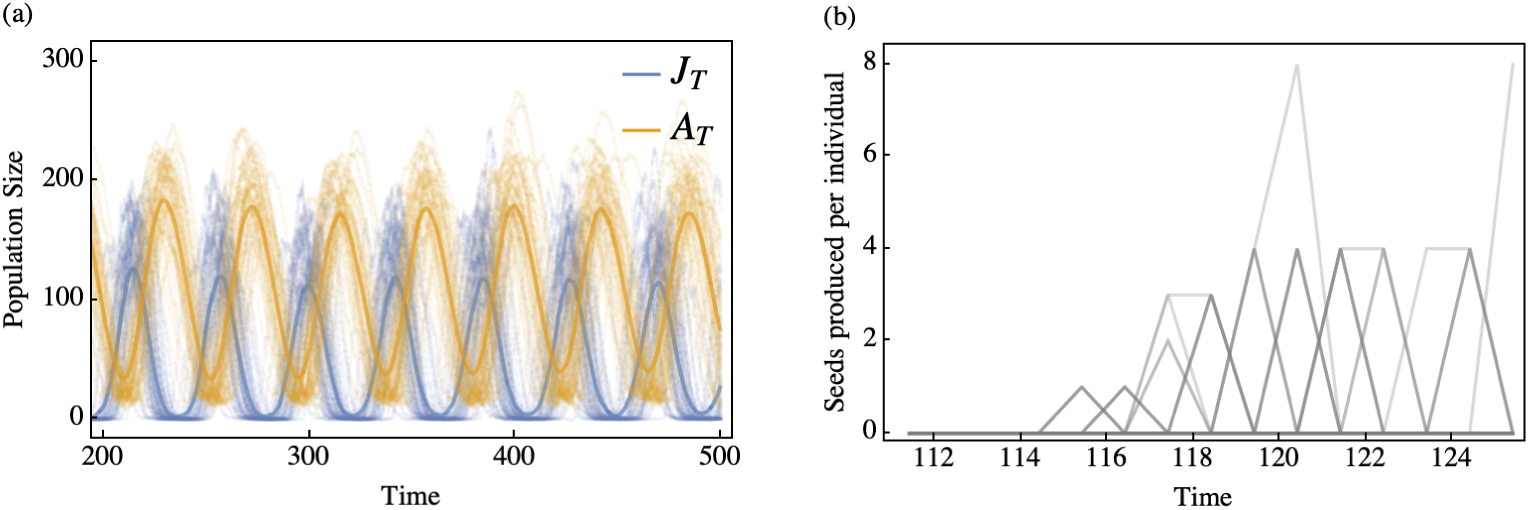
Individual-based stochastic model: population-level cycles and individual-level reproductive time series. Parameter set corresponds to case C in Fig. 4, with *L* = 5, *J_T_* (0) = 50, *A_T_* (0) = 50. (a) Total juvenile (*J_T_*) and adult (*A_T_*) abundances from multiple stochastic realisations; faint and bold lines show individual realisations and their mean, respectively. (b) Seed-production time series of reproducing adults within the post-burn-in observation window, showing heterogeneous but temporally clustered reproductive output.

## 4 Masting Data

Here, we validate the resource-budget and stage-structured models against empirically observed masting patterns reported in the literature. This included time series for annual reproduction for individual plant species: *Quercus ilex*, *Quercus humilis*, *Sorbus aucuparia*, *Chionochloa pallens*, and *Fagus sylvatica* [Bogdziewicz et al., 2019, Kelly et al., 2025, Hacket-Pain et al., 2025, Bogdziewicz et al., 2024a]. To complement these individual datasets, we used data from the MASTREE + dataset [Hacket-Pain et al., 2022]. Importantly, MASTREE + records do not include reproductive output of individual trees but rather reproductive output at larger geographic scales (e.g., stands, patches, etc.). Restricting ourselves to stands, which are the smallest spatial unit, records are typically obtained from only a few individuals (traps) per stand. As such, we use the time series per stand as a proxy for individual reproductive output and consider emergent patterns of synchrony and periodicity among stands. As discussed in A, the differing scales of the individual and MASTREE + data reflect differing measures of volatility (*V̄_x_* vs. *V̄_x̄_*), synchrony (*S_x_* vs. *S_x̄_*), and periodicity (*P_x̄_* vs. *P_x̂_*).

For the MASTREE + dataset, we retain only datasets with more than one time series, as this is required to quantify synchrony and periodicity. Records with missing years or disjoint segments are excluded to avoid spurious estimates of periodicity or volatility. For multi-site studies, we restrict analyses to overlapping observation years to ensure comparability when calculating synchrony. In addition, we exclude records without a defined measurement unit to avoid conflating incompatible scales of reproductive effort. The retained studies span a broad range of units, including cones per branch or individual, cones per square meter, flowers or fruits per individual, seeds per branch or individual, biomass-based measures (g/m^2^, grains/m^3^, kg), and the percentage of individuals fruiting or reproducing. After filtering, we retained 68 stand-level studies across 37 species (Table 3 of the Supplementary Material). Regardless of the dataset/metric used, the individual-level and stand-level analyses span similar ranges in reproductive behaviour (Fig. 6), with results from the MASTREE + data varying little with the number of individuals monitored per stand.

**Fig. 6:**
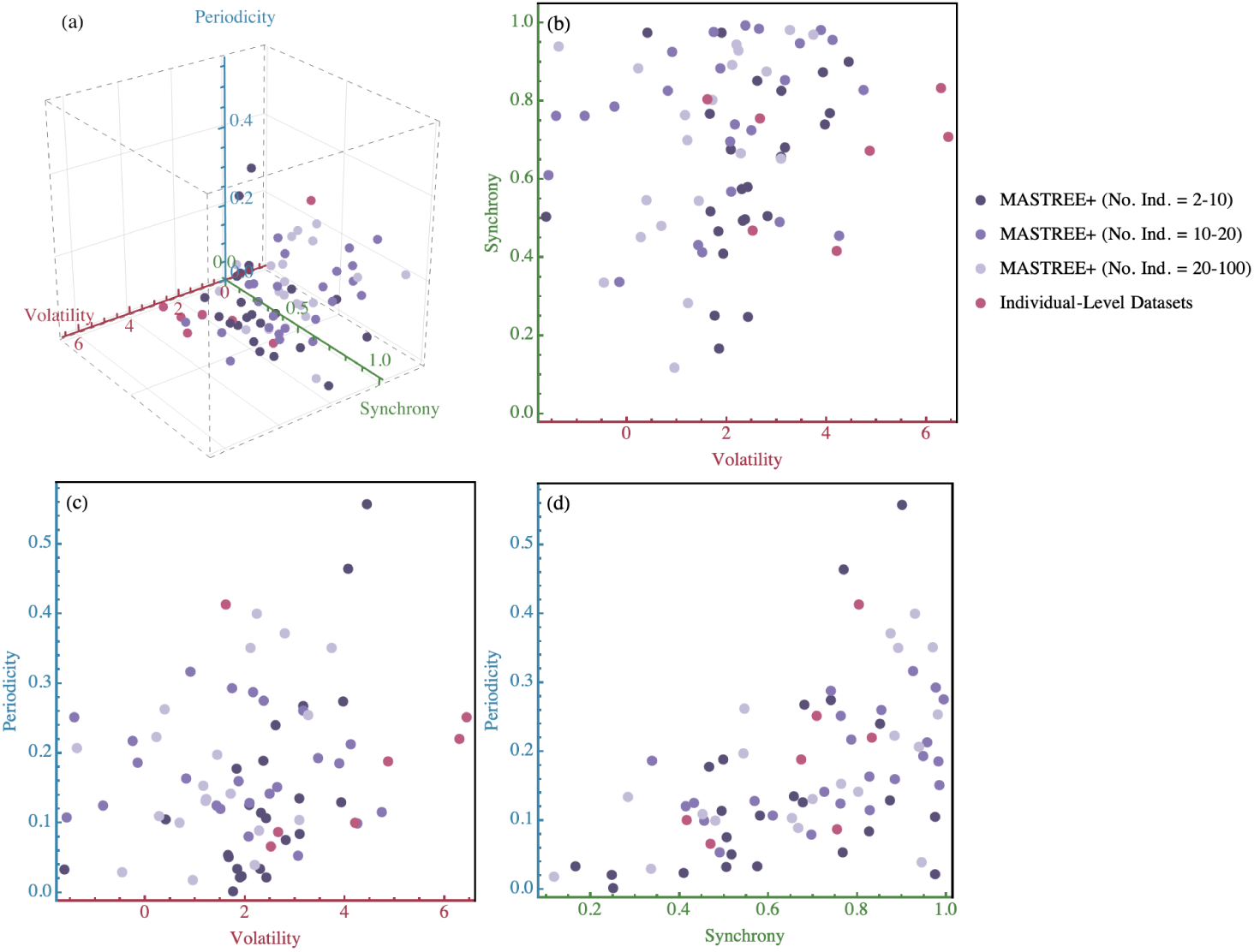
3D masting metric space for data. (a) 3D space defined by volatility, synchrony, and periodicity for the masting data. MASTREE + points correspond to stand-scale time series and are grouped according to the reported number of monitored individuals. Individual-level datasets are shown in pink for comparison. (b–d) 2D projections of the same metric space.

## 5 Model-Data Comparison

### 5.1 Sampling Model Space

For the resource-budget model, parameter sets are generated using Latin hypercube sampling (LHS) over the ranges given in Table 1, which span [0.5, 2] times the reference parameter values for each parameter. Because not all sampled combinations yield biologically admissible dynamics, we apply a rejection step to retain only parameter sets that satisfy the structural constraints of the resource-budget model: the reserve threshold must exceed the past seed production (*L_T_ > P_s_*), the cost of flowering must not exceed the reserve threshold (*C_f_* ≤ *L_T_*), and the proportionality constant must be at least unity (*R_c_* ≥ 1). Sampling continues until the required number of valid replicates is obtained. Time series of fruiting cost (*C_f_*) are then simulated for each accepted parameter set, yielding a broad ensemble of reproductive effort trajectories across the admissible parameter space. For each accepted parameter set, time series of fruiting cost (*C_f_*) are simulated for 15 time units, consistent with the mean length of the empirical masting records (14 years), yielding a broad ensemble of reproductive effort trajectories across the admissible parameter space.

For the stage-structure model, the qualitative dynamics are determined entirely by the number of sub-stages (*p*), the juvenile crowding weight (*ɛ*), and maximum seed set (*L*); all remaining parameters are held fixed at biologically plausible values reflecting perennial plant life histories (Table 2). We sample combinations of (*p, ɛ*) using Latin hypercube sampling in the same manner as for the resource-budget model, and sample *L* over the interval (0, 10000) to cover the broad range of reproductive capacities observed among perennial plants. All simulations are initialised at *J_T_* (0) = 50 total juveniles and *A_T_* (0) = 50 total adults distributed uniformly across the *p* sub-stages. Each simulation is run for *t*_max_ + 10 *t_w_* time units, where *t*_max_ = 100 is a burn-in period that avoid early transient dynamics and *t_w_* = 15 is the width of the observation window, chosen to be of the same order as the observed masting time series lengths.

The observation window is anchored at *t*_0_, the time of the first birth event after *t*_max_, and spans [*t*_0_*, t*_0_ + *t_w_*). Within this window we retain reproductive time series from all adults who produced at least one offspring in [*t*_0_*, t*_0_ +*t_w_*) and construct individual-level seed-production time series by binning birth events into non-overlapping unit intervals, yielding a regularly-spaced series of length *t_w_* = 15 per individual. Simulation runs in which no birth event occurs within [*t*_max_*, t*_max_ + 10 *t_w_*) = [100, 250) are excluded from the analysis.

### 5.2 Metric-Space Comparison

To compare the stage-structured and resource-budget mechanisms with empirical masting patterns, we place model outputs and data in a common metric framework based on the continuous, spectral, and correlation-based measures summarized by Qiu et al. [2023]. Specifically, we evaluate replicate time series using volatility, synchrony, and periodicity, which quantify complementary features of reproductive dynamics.

Observed masting time-series (MASTREE + subset and inidividual-level datasets) are compared with those generated by mechanistic stage-structured and resource-budget models within a three-dimensional masting metrics space, as shown in Fig. 7. Both the stage-structured and resource-budget models partially overlap with the empirical data cloud across the three masting metrics. However, these models occupy distinct regions within the three-dimensional volatility, synchrony, and periodicity space, indicating that each captures different dynamical regimes of masting behavior. The degree of overlap with the observed data varies across metric pairs. Along the periodicity axis, both models span nearly the full range observed in the empirical datasets (Fig. 7c and 7d). In contrast, both models are concentrated at relatively low volatility values and do not reproduce the high-volatility tail present in many empirical time series (Fig. 7b and 7c). Along the synchrony axis, the resource-budget model occupies a narrower domain, clustering at high synchrony values, whereas the stage-structured model is more broadly distributed and overlaps more extensively with the empirical data cloud (Fig. 7b and 7d). The centroid of the stage-structured model lies close to the empirical distribution in volatility and synchrony space, indicating that demographic stochasticity within a stage-structured framework can reproduce realistic joint patterns of these two metrics as observed in natural masting systems (Fig. 7b). Furthermore, the stage-structured model exhibits correlated variation across all three masting metrics simultaneously. In contrast, the resource-budget model displays a tighter internal relationship primarily between volatility and periodicity, with synchrony varying more independently of the other two metrics (Fig. 7b and 7d).

**Fig. 7:**
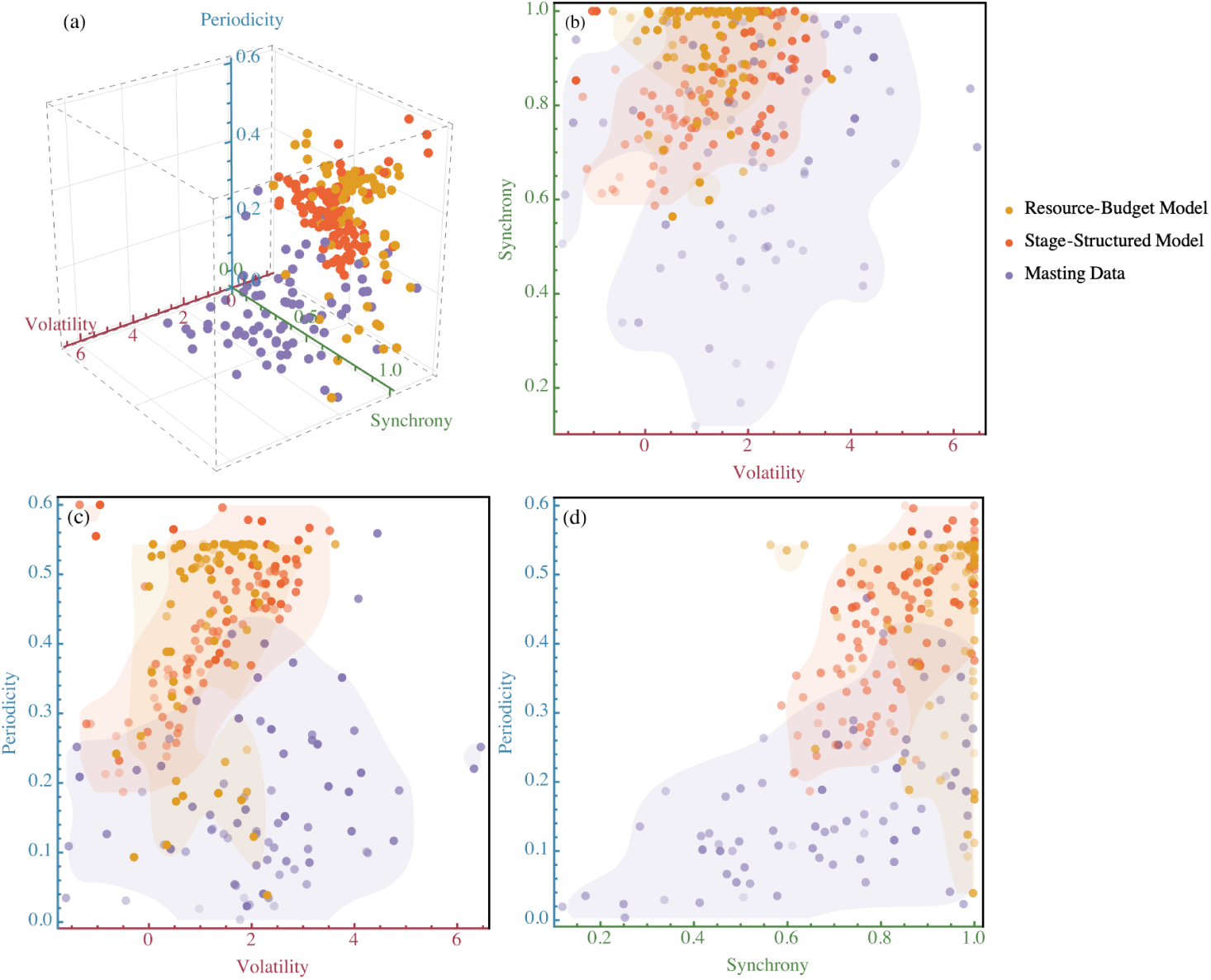
3D masting metric space for the mechanistic models and data. (a) 3D panel showing orientation of the metric space. Panels (b)-(d): 2D projects of 3D space. Opacity of points represents the intensity of the third omitted metric (darker = higher). Shaded areas show a non-parametric 95% kernel-density regions for each dataset. All axes ranges were set to span the theoretical predictions and empirical observations ±10%.

### 5.3 PCA-Based Comparison

While each metric represents a distinct aspect of reproductive dynamics, they are not mutually exclusive in practice. To more effectively characterize the structure of variation and facilitate comparison between theoretical models and observed data, principal component analysis (PCA) was applied to the masting dataset to establish an empirical reference frame. The resulting principal components (PC1–PC3) serve as fixed, datadriven axes for projecting the outputs of both models (Fig. 8). PC1 and PC2 together account for 84.26% of the variance in the empirical dataset (55.72% and 28.74%, respectively). PC3 explains an additional 15.54%, underscoring the necessity of a comprehensive three-dimensional description of masting dynamics rather than relying on a single- or two-metric summary. The loadings reveal that PC1 reflects a balanced contribution from periodicity (approximately 40%) and synchrony (approximately 38%), with a smaller contribution from volatility (approximately 15%), indicating that these metrics collectively describe core masting behaviour (Fig. 8c). PC2 is primarily influenced by volatility (approximately 80%), with minor contributions from periodicity and synchrony. In contrast, PC3 is shaped by nearly equal contributions from periodicity (approximately 50%) and synchrony (approximately 45%), with negligible loading from volatility. In the PC1–PC2 projection, both models cluster in the negative PC1 region relative to the empirical data, indicating that simulated time series tend to exhibit stronger combined periodicity and synchrony than observed masting records (Fig. 8a). The stage-structured model more closely aligns with empirical variation along PC1 and PC3, reflecting more realistic combinations of synchrony and periodicity. In contrast, the resource-budget model diverges substantially along PC2, indicating systematically higher volatility in its outputs. The kernel density distributions of PC scores further illustrate these differences: along PC1, both models shift toward more negative values than the empirical data; along PC2, the resource-budget model is strongly displaced toward negative scores; and along PC3, the two models bracket the empirical distribution from opposite sides (Fig. 8b). Overall, PCA demonstrates that the two models reproduce different modes of masting behaviour, with neither fully capturing the multivariate structure of the empirical data.

**Fig. 8:**
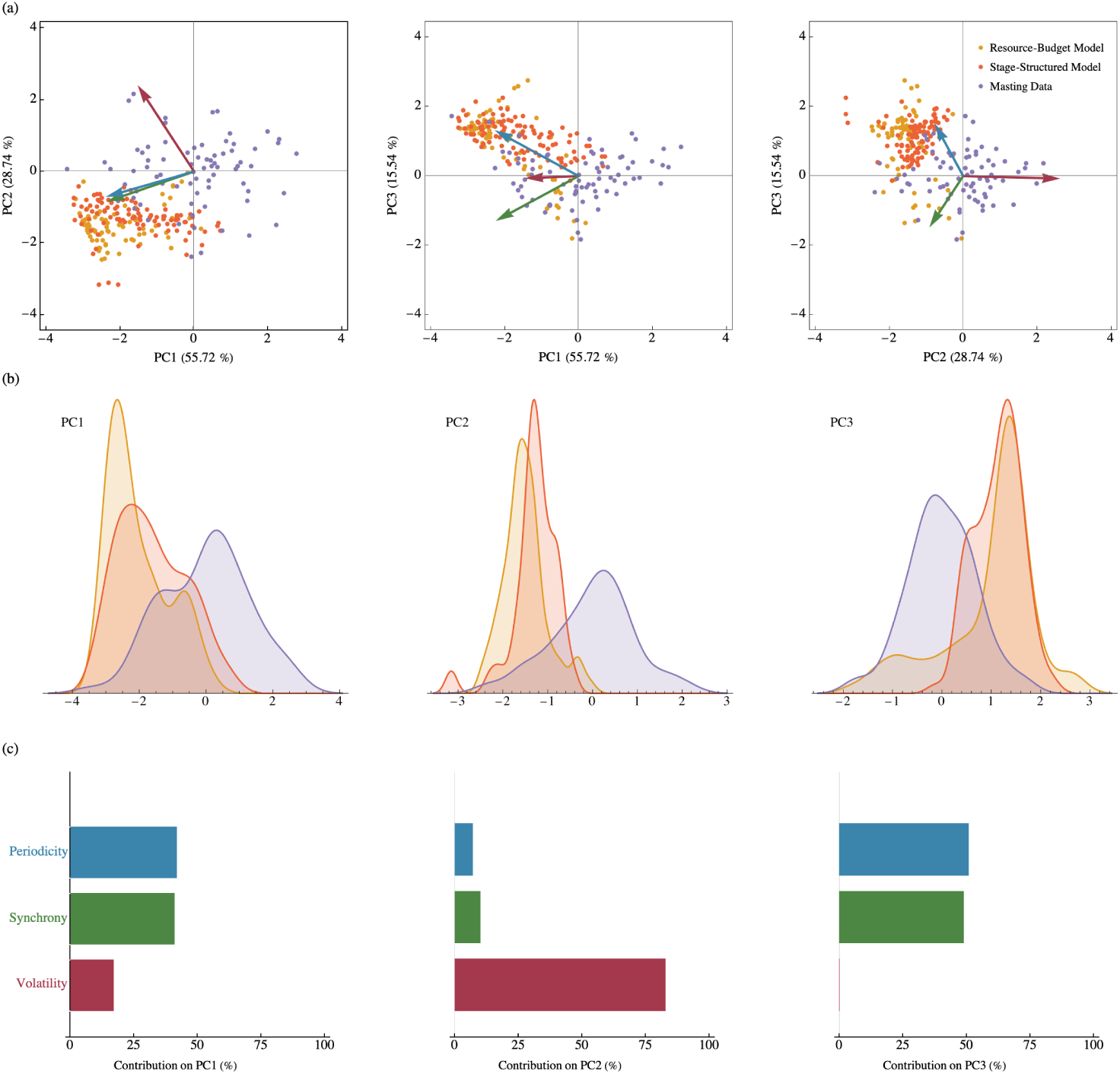
PCA of masting metrics across models and data. (a) Biplots projecting the resource-budget model, stage-structured model, and masting data onto the PC1–PC2, PC1–PC3, and PC2–PC3 planes; axis labels report the variance explained by the empirical dataset and arrow lengths reflect the magnitude of metric loadings. (b) Kernel density estimates of PC scores for each dataset along PC1, PC2, and PC3. (c) Bar charts showing the percentage contribution of periodicity, synchrony, and volatility to each principal component.

## 6 Discussion

Consistent with Shaw et al. [2024] first persona for framing theoretical work, we present our model as a novel mechanism to explain a specific empirical pattern: the highly variable, intermittent, and synchronized reproduction known as masting. In our model, life-history delays generate cyclic changes in population structure, and demographic stochasticity converts these cycles into punctuated, irregular realized reproduction at the individual level; because event hazards depend on the shared population state, these pulses can become temporally aligned, producing masting-like co-fluctuations. We emphasize that this is a baseline, non-seasonal mechanism that does not include weather cues or other explicit environmental forcing.

Our stage-structured population model represents a distinct mode of reproductive synchronization compared to the classic resource-budget framework, and highlights that other, yet to be characterized, processes may contribute to similar reproductive patterns in nature. Despite the models’ distinct biological foundations, both incorporate delays. Cyclic pattern as masting descriptor emerges from internal delays either explicitly through stage progression in our model or implicitly through resource accumulation in resource-budget model which suggests that delayed feedbacks may be a fundamental ingredient of masting dynamics. This insight connects to other aspects of population biology, where variation in the timing of life-history events such as maturation, reproduction, infection, or dormancy profoundly influence ecological, epidemiological, and evolutionary outcomes. Our findings also underscore how delays act in combination with variation in the strength of density dependence among lifehistory stages to generate collective periodic behaviour. In the stochastic setting, these same density-dependent interactions are also responsible for generating heterogeneity among individuals, as competition mediates how demographic fluctuations translate into variability in individual reproductive output. Cyclic reproduction arises in the stage-structured model when juveniles contribute only weakly to density-dependent population growth. Analogously, in the SIR epidemiological model, cyclic outbreaks occur as a result of a sustained resistant phase during which hosts can not be reinfected and hence do not contribute to mass-action transmission. Hence, in both models there exists a stage (e.g., juvenile or resistant) in which individuals are relatively inert and contribute little to population dynamics (e.g., reproduction or infection) before re-emerging in a subsequent stage (e.g., maturation or loss of immunity). Here, synchrony refers to temporal co-fluctuation among replicated reproductive time series over overlapping years, and we did not analyze spatial synchrony across locations. Whereas we focused on cue-free baseline dynamics in a single, well-mixed population, future extensions could consider how variation in competition across spatial structured environments, as well as among life-history stages, may contribute to the emergence and persistence of masting dynamics.

Beyond the role of delays and density dependence, our stage-structured framework highlights demographic stochasticity as the primary source of individual-level variability. Because reproduction, maturation, and mortality occur as discrete events, demographic stochasticity can generate intermittent individual output and some degree of temporal alignment via shared state dependence; in many systems, weather cues are likely to further synchronize and stabilize these dynamics. As in quasi-periodic epidemic outbreaks, the ability for demographic stochasticity to contribute to emergent phenomena such as synchronous reproduction may be a general principle with widespread implications for population biology. In the resource–budget model, by contrast, reproductive synchrony arises through the deterministic process of pollen limitation: when few individuals flower in a given year, pollination success is low, favoring simultaneous reproduction in subsequent years as resource reserves rebuild. Although environmental stochasticity was not included in the present analysis, a growing literature suggests that correlated weather cues commonly synchronize reproduction across individuals, while environmental effects on flowering and pollination efficiency may further amplify or dampen seed set [Ferńandez-Martínez et al., 2017]. Future work will incorporate cue-based forcing and assess its effects on synchrony and periodicity.

We have focused primarily on the proximate mechanisms of masting, but its ultimate causes are equally important. The hypothesized ultimate benefits of masting are rooted in economies of scale (EOS) which describes how increasing reproductive output reduces the relative cost per unit. Among the most widely supported EOS hypotheses for the evolution of masting are pollen efficiency [Kelly et al., 2001] and predator satiation [Janzen, 1971]. Since pollen limitation is necessary for the emergence of synchronous reproduction in the resource-budget model, to date evolutionary theory has focused on the role of pollen efficiency either in isolation or in combination with other EOS [Isagi et al., 1997, Satake and Iwasa, 2000, 2002a, Tachiki and Iwasa, 2008]. By providing an alternative proximate mechanism that, in our baseline formulation, does not assume pollen limitation, the stage-structured model offers a foundation for testing how economies of scale (including pollen efficiency and predator satiation) might act, alone or in combination, when synchrony arises from life-history timing and demographic stochasticity. Importantly, this does not preclude cue-based or pollen-mediated effects; rather, it highlights a complementary pathway that may interact with them.

Masting plays a pivotal role in linking plant reproductive strategies to ecosystem services and conservation outcomes. Because mast seeding is inherently volatile and spatially localized, its variability cascades through food webs and influences species interactions across trophic levels [Clark et al., 2019]. Recent evidence indicates that climate change is disrupting the frequency and synchrony of mast events, and thereby reshaping their ecological and evolutionary benefits [Bogdziewicz et al., 2020, Hacket-Pain and Bogdziewicz, 2021]. Given that masting drives key processes such as forest regeneration, seed predator dynamics, and disease transmission, understanding how these dynamics respond to climate change is crucial for effective conservation and land management [Pearse et al., 2021]. By capturing how masting arises from life history, density dependence and stochasticity, all of which may depend on climate, our stage-structured model offers new opportunities for predicting the effects of climate change. Importantly, understanding the proximate mechanisms underlying masting is also essential for predicting how climate change will influence this reproductive behavior. Together, the stage-structured and resource budget models provide a more general theoretical foundation for masting and open pathways for future work on prediction and ecosystem management.

## Declarations

- **Funding** This research is supported by funds from the National Science and Engineering Research Council (CRC-2021-00276 and RGPIN-2022-03113 to A.M. and RGPIN-2021-04088 to J.M.S.). M.S. is supported in part by funds from Simon Fraser University.
- **Conflict of interest/Competing interests** The authors have no conflicts of interest to declare.
- **Ethics approval and consent to participate** Not applicable
- **Consent for publication** All authors approve the submission of the manuscript.
- **Data availability** Previously published reproductive output data from MASTREE+ [Hacket-Pain et al., 2022] were utilized and are publicly available at https://github.com/JJFoest/MASTREEplus. Additional individual-level reproductive output datasets analyzed in this study are accessible through the data-availability statements and repositories associated with the cited studies [Bogdziewicz et al., 2019, Kelly et al., 2025, Hacket-Pain et al., 2025, Bogdziewicz et al., 2024a].
- **Materials availability** Not applicable
- **Code availability** The accompanying *Mathematica* code used for data processing and computation of masting metrics is provided with the submission and will be archived on Zenodo upon manuscript acceptance.
- **Author contribution** M.S. developed the model, conducted all analyses, and drafted the manuscript. A.M. led the planning and supervision of the research and provided critical revisions. J.M.S. contributed to supervision and manuscript editing.

## Appendix A Masting Metrics

Masting is challenging to quantify. Numerous metrics have been used throughout the literature to capture characteristic features of this reproductive behaviour (see Table 2 of the Supplementary Material), but these metrics are rarely used in combination (but see Qiu et al. [2023]). We categorize the available metrics into three groups based on the primary feature of masting that they capture. These include: 1) inter-annual variation in the seed set of an individual tree, 2) synchrony in the reproductive effort of trees within a population or stand, and 3) the emergent periodicity in seed production at the population level. It is important to note that while these features are distinct from one another, they are interdependent; for example, individual-level variation is necessary for periodicity at the population-wide scale. Here, we adopt the unifying framework of spectral and correlation-based metrics recently summarized by Qiu et al. [2023] with their implementation provided in the accompanying *Mathematica* file.

Let *x*(*t*) be the reproductive effort of an individual at discrete time *t* ∈ {*k*_1_*, k*_1_ + 1*, . . ., k*_2_} where *n* = *k*_2_ − *k*_1_ is the length of the observed time series. The population-level reproductive output can then be quantified as the average reproductive effort across individuals, which we denote by *x̄*(*t*) = E*_i_*[*x_i_*(*t*)]. And the mean reproductive output among populations (stands) is denoted as *x̂* (*t*) = E*_p_*[*x̄*(*t*)]. For discrete time series *x*(*t*), patterns in the temporal variation can be expressed in the frequency domain, *ω*, via the spectral density

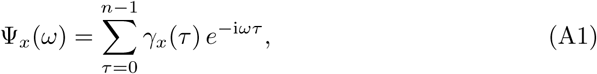

where *γ_x_*(*τ*) is the auto-covariance function of *x*(*t*) at lag *τ* ∈ {0, 1, 2, · · · *, N* − 1}. Conversely, the auto-covariance function can be recovered from the spectral density via an inverse discrete Fourier transform:

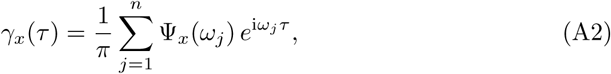

where *ω_j_* = 2*πj/n* for *j* ∈ {1, 2, · · · *, N* }. Notably, at zero lag, *γ_x_*(0) equals the total variance of the time series, also known as spectral variance,

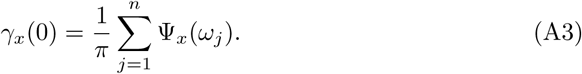

To capture the first defining feature of masting, we follow Qiu et al. [2023] and define individual volatility, which we denote as V*_x_* (note that our notation differs slightly from that of Qiu et al. [2023]), as a standardized and period-weighted spectral variance,

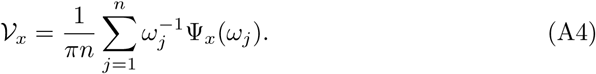

This period-weighted average captures variation in individual reproductive effort concentrated at low frequencies (i.e., over longer time scales). The average volatility among individuals in the population is then given by:

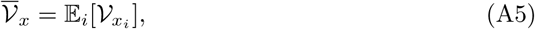

which we use as our focal metric for volatility in the mechanistic models and in the individual-level datasets; for the stand-level MASTREE + records, which we use as a proxy for individual-level dynamics, we report the analogous quantity *V̄_x_* instead. Since volatility can vary widely, we consider its log value. The volatility metric is well-suited to detecting the hallmark of masting, specifically large swings in reproductive effort that occur over many years rather than from one year to the next. This makes volatility more informative than traditional measures such as the coefficient of variation, which may underrepresent the contribution of multi-year cycles. We can also calculate an analogous measure of volatility in population-level reproductive output, denoted *V̄_x_*, which is an emergent feature of individual-level variation, but in our analysis, we do not consider this as a unique feature of masting (note *V̄_x̄_* ≠ *V̄_x̄_*).

We next consider synchrony in reproductive effort among the individuals in the population. Again following Qiu et al. [2023] in spirit but with distinct notation

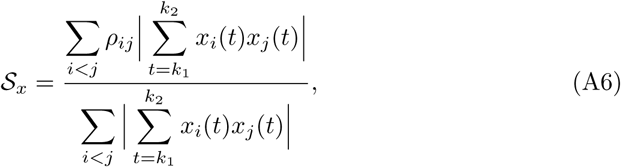

where *ρ_ij_* denotes the pairwise correlation between reproductive effort of individual *i*, *x_i_*(*t*), and individual *j*, *x_j_*(*t*). Unlike simple correlation-based measures, S*_x_* weights correlations by the magnitude of reproductive output, thereby emphasizing synchrony in mast years when ecological impacts are greatest. Although computationally intensive and less standardized in the literature, S*_x_* provides a powerful measure of the collective reproductive dynamics in a population. We use this as our focal synchrony metric for the mechanistic models and the individual-level datasets, whereas for the stand-level MASTREE + records we use the corresponding stand-level quantity S*_x_*.

Finally, we turn to population-level periodicity, which measures the extent to which variation in reproductive output is concentrated at low frequencies,

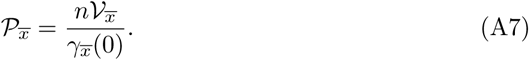

As with *V̄_x_* at the individual scale, *P_x_* measures the extent to which a population’s reproductive output is dominated by multi-year patterns rather than annual fluctuations. In masting studies,this definition of periodicity helps distinguish between species that show irregular but strongly cyclical reproduction versus those with more diffuse, short-term variation. Just as we may calculate population-level volatility, we can also calculate individual-level periodicity *P_x̄_* = *nV_x_/γ_x_*(0), but we focus only on population-level periodicity as a characteristic of masting; accordingly, we use P*_x_* for the mechanistic models and the individual-level datasets, while for the stand-level MASTREE + records we use the analogous across-stand quantity *P_x̂_*.

## Appendix B Floquet Theory

To assess the stability of sustained cyclic dynamics observed in our stage-structured model for certain parameter combinations of (*p, ɛ*) (see Fig. 4), we applied Floquet theory. This approach analyzes small perturbations around a periodic solution of the system and determines whether they grow or decay over time. Floquet theory has been widely used in theoretical ecology to study the stability of periodic solutions (e.g., Klausmeier [2008]), making it a natural choice for our analysis.

First, we numerically computed a limit cycle solution by integrating the system of ordinary differential equations over one period *T*, using periodicity conditions as constraints to ensure that the sub-stages (*J_i_* and *A_j_*) at *t* = 0 match those at *t* = *T* . The resulting solution defines a closed orbit in the solution space representing sustained cyclic dynamics. Next, we constructed the time-dependent Jacobian matrix

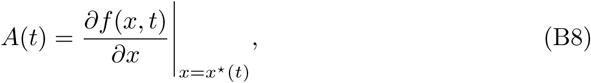

evaluated along the limit cycle trajectory *x*\*(*t*). We then solved the following equation

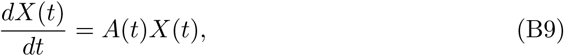

given initial condition *X*(0) = *I*, over a single period *T* to obtain the monodromy matrix *X*(*T*). The eigenvalues of this matrix, called Floquet multipliers, determine the local stability of the cycle: if all multipliers have real parts less than one in absolute value (except the trivial multiplier corresponding to time translation), the limit cycle is stable. Finally, we summarized the stability properties across parameter space by computing the largest real part of the Floquet multipliers for each parameter pair (*p, ɛ*). Parameter combinations yielding max |Re(*λ*)| *<* 1 correspond to stable cycles, whereas max |Re(*λ*)| *>* 1 indicate instability and the onset of further bifurcations.

## Appendix C Individual-Based Stochastic Process

Our implementation follows a two-step procedure provided as pseudo-code in Algorithm 1. First, we determine which type of event occurs next based on the total rates of births, aging, and deaths. Then, having chosen an event type, we randomly select which stage experiences that event, with probability weighted by current stage abundances (where applicable). After the state is updated (including sampling Δ*J*_1_ for reproduction), we recompute all event rates and repeat until the simulation time reaches the maximum horizon or no further events are possible. This approach exactly reproduces the dynamics of a fully enumerated Gillespie simulation while keeping the event list compact and the simulation efficient.

**Algorithm 1.**
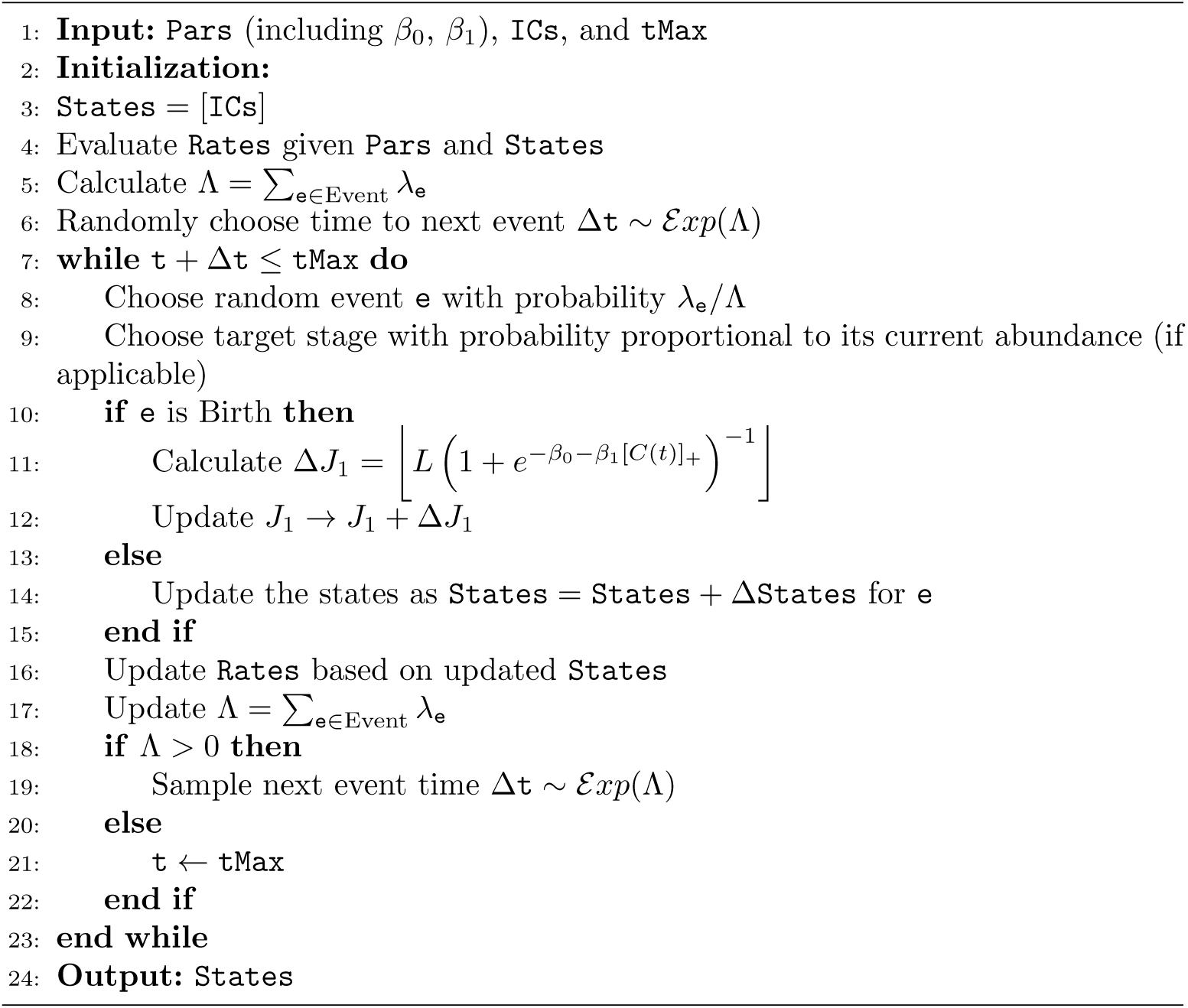
Gillespie algorithm with two-level sampling for the stage-structured model. The first level chooses an event type; the second level selects a target stage (where applicable). For birth events, the state update is stochastic.

## References

1. J. J. Beck, M. J. McKone, and S. Wagenius. Masting, fire-stimulated flowering, and the evolutionary ecology of synchronized reproduction. Ecology, 105(4):e4261, 2024. doi: 10.1002/ecy.4261.

2. M. Bogdziewicz, M. Żywiec, J. M. Espelta, et al. Environmental veto synchronizes mast seeding in four contrasting tree species. The American Naturalist, 194(2): 246–259, 2019. doi: 10.1086/704111.

3. M. Bogdziewicz, D. Kelly, A. J. Tanentzap, P. A. Thomas, J. G. Lageard, and A. Hacket-Pain. Climate change strengthens selection for mast seeding in European beech. Current Biology, 30(17):3477–3483.e2, 2020. doi: 10.1016/j.cub.2020.06.056.

4. M. Bogdziewicz, I. Chybicki, J. Szymkowiak, B. Ulaszewski, J. Burczyk, et al. Relatives reproduce in synchrony: Kinship and individual condition shape intraspecific variation in masting phenotype. Proceedings of the Royal Society B: Biological Sciences, 291(2017):20232732, 2024a. doi: 10.1098/rspb.2023.2732.

5. M. Bogdziewicz, D. Kelly, D. Ascoli, T. Caignard, F. Chianucci, E. E. Crone, E. Fleurot, J. J. Foest, G. Gratzer, T. Hagiwara, Q. Han, V. Jourńe, L. Keurinck, K. Kondrat, R. McClory, J. M. LaMontagne, I. A. Mundo, A. Nussbaumer, I. Oberklammer, M. Ohno, I. S. Pearse, M. B. Pesendorfer, G. Resente, A. Satake, M. Shibata, R. S. Snell, J. Szymkowiak, L. Touzot, R. Zwolak, M. Zywiec, and A. J. Hacket-Pain. Evolutionary ecology of masting: mechanisms, models, and climate change. Trends in Ecology & Evolution, 39(9):851–862, 2024b. doi: 10.1016/j.tree.2024.05.006.

6. C. Bregnard, O. Rais, and M. J. Voordouw. Masting by beech trees predicts the risk of Lyme disease. Parasites & Vectors, 14(1):168, 2021. doi: 10.1186/s13071-021-04646-0.

7. J. S. Clark, C. L. Nuñez, and B. Tomasek. Foodwebs based on unreliable foundations: spatiotemporal masting merged with consumer movement, storage, and diet. Ecological Monographs, 89(4):e01381, 2019. doi: 10.1002/ecm.1381.

8. E. E. Crone, E. J. B. McIntire, and J. Brodie. What defines mast seeding? Spatiotemporal patterns of cone production by whitebark pine. Journal of Ecology, 99(2): 438–444, 2011. doi: 10.1111/j.1365-2745.2010.01790.x.

9. M. Ferńandez-Martínez, S. Vicca, I. A. Janssens, J. M. Espelta, and J. Peñuelas. The North Atlantic Oscillation synchronises fruit production in western European forests. Ecography, 40(7):864–874, 2017. doi: 10.1111/ecog.02296.

10. J. J. Foest, M. Bogdziewicz, M. B. Pesendorfer, D. Ascoli, A. Cutini, A. Nussbaumer, A. Verstraeten, B. Beudert, F. Chianucci, F. Mezzavilla, G. Gratzer, G. Kunstler, H. Meesenburg, M. Wagner, M. Mund, N. Cools, S. Vacek, W. Schmidt, Z. Vacek, and A. Hacket-Pain. Widespread breakdown in masting in European beech due to rising summer temperatures. Global Change Biology, 30(5):e17307, 2024. doi: 10.1111/gcb.17307.

11. D. T. Gillespie. A general method for numerically simulating the stochastic time evolution of coupled chemical reactions. Journal of Computational Physics, 22(4): 403–434, 1976. doi: 10.1016/0021-9991(76)90041-3.

12. S. Goņcalves, G. Abramson, and M. F. C. Gomes. Oscillations in SIRS model with distributed delays. The European Physical Journal B, 81(3):363–371, 2011. doi: 10.1140/epjb/e2011-20054-9.

13. A. Hacket-Pain and M. Bogdziewicz. Climate change and plant reproduction: trends and drivers of mast seeding change. Philosophical Transactions of the Royal Society B: Biological Sciences, 376(1839):20200379, 2021. doi: 10.1098/rstb.2020.0379.

14. A. Hacket-Pain, J. Szymkowiak, V. Jourńe, M. K. Barczyk, P. A. Thomas, J. G. A. Lageard, D. Kelly, and M. Bogdziewicz. Growth decline in European beech associated with temperature-driven increase in reproductive allocation. Proceedings of the National Academy of Sciences, 122(5):e2423181122, 2025. doi: 10.1073/pnas.2423181122.

15. A. Hacket-Pain, J. J. Foest, I. S. Pearse, J. M. LaMontagne, W. D. Koenig, G. Vacchiano, M. Bogdziewicz, et al. Mastree+: Time-series of plant reproductive effort from six continents. Global Change Biology, 28(9):3066–3082, 2022. doi: 10.1111/gcb.16130.

16. H. W. Hethcote. The mathematics of infectious diseases. SIAM Review, 42(4):599–653, 2000. doi: 10.1137/S0036144500371907.

17. P. J. Hurtado and A. S. Kirosingh. Generalizations of the ‘Linear Chain Trick’: incorporating more flexible dwell time distributions into mean field ODE models. Journal of Mathematical Biology, 79(5):1831–1883, 2019. doi: 10.1007/s00285-019-01412-w.

18. Y. Isagi, K. Sugimura, A. Sumida, and H. Ito. How does masting happen and synchronize? Journal of Theoretical Biology, 187(2):231–239, 1997. doi: 10.1006/jtbi.1997.0442.

19. Y. Ito and G. Kudo. The contribution of carbon budget to masting intervals in *Veratrum album* populations inhabiting different elevations. American Journal of Botany, 111(3):e16295, 2024. doi: 10.1002/ajb2.16295.

20. D. H. Janzen. Seed predation by animals. Annual Review of Ecology, Evolution, and Systematics, 2:465–492, 1971. doi: 10.1146/annurev.es.02.110171.002341.

21. V. Jourńe, A. Hacket-Pain, I. Oberklammer, M. B. Pesendorfer, and M. Bogdziewicz. Forecasting seed production in perennial plants: identifying challenges and charting a path forward. New Phytologist, 239(2):466–476, 2023. doi: 10.1111/nph.18957.

22. D. Kelly. The evolutionary ecology of mast seeding. Trends in Ecology & Evolution, 9(12):465–470, 1994. doi: 10.1016/0169-5347(94)90310-7.

23. D. Kelly, D. E. Hart, and R. B. Allen. Evaluating the wind pollination benefits of mast seeding. Ecology, 82(1):117–126, 2001. doi: 10.1890/0012-9658(2001)082[0117:ETWPBO]2.0.CO;2.

24. D. Kelly, J. Szymkowiak, A. Hacket-Pain, and M. Bogdziewicz. Fine-tuning mast seeding: as resources accumulate, plants become more sensitive to weather cues. New Phytologist, 246(5):1975–1985, 2025. doi: 10.1111/nph.70092.

25. C. A. Klausmeier. Floquet theory: a useful tool for understanding nonequilibrium dynamics. Theoretical Ecology, 1(3):153–161, 2008. doi: 10.1007/s12080-008-0016-2.

26. J. D. Lauder, E. V. Moran, and S. C. Hart. Fight or flight? Potential tradeoffs between drought defense and reproduction in conifers. Tree Physiology, 39(7):1071–1085, 2019. doi: 10.1093/treephys/tpz031.

27. A. L. Lloyd. Destabilization of epidemic models with the inclusion of realistic distributions of infectious periods. Proceedings of the Royal Society of London B: Biological Sciences, 268(1470):985–993, 2001. doi: 10.1098/rspb.2001.1599.

28. A. MacPherson and S. P. Otto. Joint coevolutionary–epidemiological models dampen Red Queen cycles and alter conditions for epidemics. Theoretical Population Biology, 122:137–148, 2018. doi: 10.1016/j.tpb.2017.12.003.

29. M. Nuño, Z. Feng, M. Martcheva, and C. Castillo-Chavez. Dynamics of two-strain influenza with isolation and partial cross-immunity. SIAM Journal on Applied Mathematics, 65(3):964–982, 2005. doi: 10.1137/S003613990343882X.

30. I. S. Pearse, W. D. Koenig, and D. Kelly. Mechanisms of mast seeding: resources, weather, cues, and selection. New Phytologist, 212(3):546–562, 2016. doi: 10.1111/nph.14114.

31. I. S. Pearse, J. M. LaMontagne, M. Lordon, A. L. Hipp, and W. D. Koenig. Biogeography and phylogeny of masting: do global patterns fit functional hypotheses? New Phytologist, 227(5):1557–1567, 2020. doi: 10.1111/nph.16617.

32. I. S. Pearse, A. P. Wion, A. D. Gonzalez, and M. B. Pesendorfer. Understanding mast seeding for conservation and land management. Philosophical Transactions of the Royal Society B: Biological Sciences, 376(1839):20200383, 2021. doi: 10.1098/rstb.2020.0383.

33. M. B. Pesendorfer, D. Ascoli, M. Bogdziewicz, et al. The ecology and evolution of synchronized reproduction in long-lived plants. Philosophical Transactions of the Royal Society B: Biological Sciences, 376(1839):20200369, 2021. doi: 10.1098/rstb.2020.0369.

34. T. Qiu, M.-C. Aravena Acuña, D. Ascoli, Y. Bergeron, M. Bogdziewicz, et al. Masting is uncommon in trees that depend on mutualist dispersers in the context of global climate and fertility gradients. Nature Plants, 9:1044–1056, 2023. doi: 10.1038/s41477-023-01446-5.

35. A. Satake and Y. Iwasa. Pollen coupling of forest trees: Forming synchronized and periodic reproduction out of chaos. Journal of Theoretical Biology, 203(2):63–84, 2000. doi: 10.1006/jtbi.1999.1066.

36. A. Satake and Y. Iwasa. Spatially limited pollen exchange and a long-range synchronization of trees. Ecology, 83(4):993–1005, 2002a. doi: 10.1890/0012-9658(2002)083[0993:SLPEAA]2.0.CO;2.

37. A. Satake and Y. Iwasa. The synchronized and intermittent reproduction of forest trees is mediated by the Moran effect, only in association with pollen coupling. Journal of Ecology, 90(5):830–838, 2002b. doi: 10.1046/j.1365-2745.2002.00721.x.

38. EÉ. Schermer, M.-C. Bel-Venner, D. Fouchet, A. Siberchicot, V. Boulanger, T. Caignard, M. Thibaudon, G. Oliver, M. Nicolas, J.-M. Gaillard, S. Delzon, and S. Venner. Pollen limitation as a main driver of fruiting dynamics in oak populations. Ecology Letters, 22(1):98–107, 2019. doi: 10.1111/ele.13171.

39. A. K. Shaw, A. T. Bisesi, C. Wojan, D. Kim, M. Torstenson, Naven Narayanan, P. Lutz, R. Ales, and C. Shao. Six personas to adopt when framing theoretical research questions in biology. Proceedings of the Royal Society B: Biological Sciences, 291(2031):20240803, 2024. doi: 10.1098/rspb.2024.0803.

40. Y. Tachiki and Y. Iwasa. Role of gap dynamics in the evolution of masting of trees. Evolutionary Ecology Research, 10:893–905, 10 2008.

41. Y. Tachiki and Y. Iwasa. Both seedling banks and specialist seed predators promote the evolution of synchronized and intermittent reproduction (masting) in trees. Journal of Ecology, 98(6):1398–1408, 2010. doi: 10.1111/j.1365-2745.2010.01729.x.

42. Y. Tachiki and Y. Iwasa. Evolutionary jumping and breakthrough in tree masting evolution. Theoretical Population Biology, 81(1):20–31, 2012. doi: 10.1016/j.tpb.2011.10.004.

43. Y. Tachiki and Y. Iwasa. Coevolution of mast seeding in trees and extended diapause of seed predators. Journal of Theoretical Biology, 339:129–139, 2013. doi: 10.1016/j.jtbi.2013.05.026.

44. S. Venner, A. Siberchicot, P.-F. Pélisson, E. Schermer, et al. Fruiting strategies of perennial plants: A resource budget model to couple mast seeding to pollination efficiency and resource allocation strategies. The American Naturalist, 188(1):66–75, 2016. doi: 10.1086/686684.

45. B. Zuckerberg, C. Strong, J. M. LaMontagne, S. St. George, et al. Climate dipoles as continental drivers of plant and animal populations. Trends in Ecology & Evolution, 35(5):440–453, 2020. doi: 10.1016/j.tree.2020.01.010.

