## Supplementary Materials for "An ecological model of masting reproduction matches empirical dynamics"

Table 1: Stochastic events and rates in the stage-structured model.

| Event ($\mathtt{e}$) | State change ($\Delta\mathtt{States}$) | Rate ($\lambda_{\mathtt{e}}$) | Count |
|---|---|---|---|
| $J_1$ Birth | $J_1 \to J_1 + \Delta J_1$ | $b\,[C(t)]_+\,A_T$ | 1 |
| $J_i$ Aging ($1 \le i < p$) | $J_i \to J_i - 1,\ J_{i+1} \to J_{i+1} + 1$ | $p\rho\,J_i$ | $p-1$ |
| $J_i$ Death ($1 \le i \le p$) | $J_i \to J_i - 1$ | $\delta\,J_i$ | $p$ |
| $J_p \to A_1$ | $J_p \to J_p - 1,\ A_1 \to A_1 + 1$ | $p\rho\,J_p$ | 1 |
| $A_j$ Aging ($1 \le j < p$) | $A_j \to A_j - 1,\ A_{j+1} \to A_{j+1} + 1$ | $p\nu\,A_j$ | $p-1$ |
| $A_j$ Death ($1 \le j \le p$) | $A_j \to A_j - 1$ | $\mu\,A_j$ | $p$ |
| $A_p$ Exit | $A_p \to A_p - 1$ | $p\nu\,A_p$ | 1 |

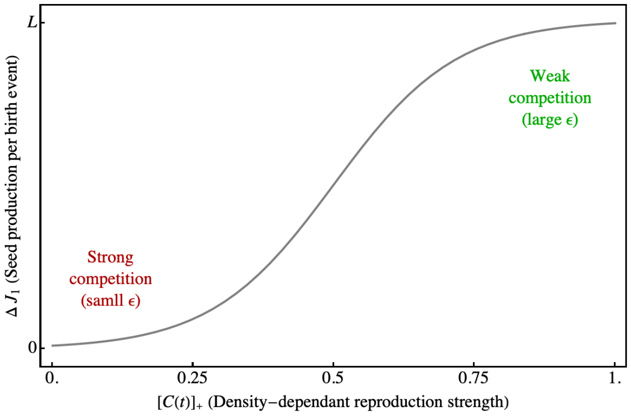


Figure 1: **Sigmoid link between the density-dependent birth strength $C(t)$ and the seed produced per reproduction event.** The parameters $\beta_0 = -4.6$ and $\beta_1 = 9.2$ are chosen so that seed set is close to $0.01L$ under strong competition and close to $0.99L$ under weak competition.

Table 2: **Summary of variation, synchrony, and periodicity metrics used in the literature.** Here, $x_i(t)$ denotes the discrete time series of reproductive output for individual $i$ at time $t$, $\bar{x}(t) = \mathbb{E}_i[x_i(t)]$ is the population mean at time $t$, and $\bar{\bar{x}} = \mathbb{E}_t[\bar{x}(t)]$ is the long-term temporal mean. Expectations $\mathbb{E}_t[\cdot]$ are taken over time, and $\mathbb{E}_i[\cdot]$ over individuals unless otherwise stated.

| Features | Metrics | Derivation |
|---|---|---|
| Volatility | Total Variance | $\mathbb{E}_t[(\bar{x}(t) - \bar{\bar{x}}(t))^2] + \mathbb{E}_t[(\bar{x}(t) - \mathbb{E}_t[\bar{x}(t)])^2]$ (Satake and Iwasa, 2002) |
| | Coefficient of Variation | $\dfrac{\sqrt{\mathrm{Var}_t[x_i(t)]}}{\mathbb{E}_t[x_i(t)]}$ (Crone et al., 2011) |
| Synchrony | Mean Pearson Correlation | $\dfrac{1}{\binom{N}{2}} \sum_{i<j} \dfrac{\mathrm{Cov}_t[x_i(t), x_j(t)]}{\sqrt{\mathrm{Var}_t[x_i(t)]\mathrm{Var}_t[x_j(t)]}}$ (Bogdziewicz et al., 2019) |
| | Spearman Rank Correlation | $\dfrac{1}{\binom{N}{2}} \sum_{i<j} \dfrac{\mathrm{Cov}_t[R_i(t), R_j(t)]}{\sqrt{\mathrm{Var}_t[R_i(t)]\mathrm{Var}_t[R_j(t)]}}$ (LaMontagne et al., 2024) |
| | Global Synchrony | $\dfrac{\mathbb{E}_t[(\bar{x}(t) - \bar{\bar{x}}(t))^2]}{\mathbb{E}_t[(\bar{x}(t) - \bar{\bar{x}}(t))^2] + \mathbb{E}_t[\mathbb{E}_i[(x_i(t) - \bar{x}(t))^2]]}$ (Satake and Iwasa, 2002) |
| | Standardized Synchrony | $\dfrac{\mathrm{Var}_t[\mathbb{E}_i[x_i(t)]]}{\left(\sum_{i=1}^{n} \sigma_{x_i}\right)^2}$ (Loreau and De Mazancourt, 2008) |
| Periodicity | Autocorrelation | $\rho(k) = \dfrac{\mathbb{E}_t[(\bar{x}(t) - \mathbb{E}_t[\bar{x}(t)])(\bar{x}(t+k) - \mathbb{E}_t[\bar{x}(t)])]}{\mathbb{E}_t[(\bar{x}(t) - \mathbb{E}_t[\bar{x}(t)])^2]}$ (Woodward et al., 2022) |

Table 3: MASTREE+ subset used in this study.

| Species | Study ID | # individuals | Unit | Species | Study ID | # individuals | Unit |
|---|---|---|---|---|---|---|---|
| *Quercus spp.* | P05128 | 20 | seeds/$m^2$ | *Fagus crenata* | M00012 | 10 | seeds/$m^2$ |
| *Fagus crenata* | M04024 | 5 | seeds/$m^2$ | *Fagus crenata* | P05163 | 10 | seeds/$m^2$ |
| *Quercus rubra* | P05035 | 90 | seeds/$m^2$ | *Picea abies* | D00035 | 81 | seeds/$m^2$ |
| *Fagus sylvatica* | A00120 | 10 | seeds/$m^2$ | *Fagus sylvatica* | D00035 | 81 | seeds/$m^2$ |
| *Picea engelmannii* | M00284 | 10 | seeds/$m^2$ | *Abies alba* | D06205 | 9 | seeds/$m^2$ |
| *Abies alba* | D00035 | 81 | seeds/$m^2$ | *Quercus petraea* | D00036 | 10 | seeds/$m^2$ |
| *Pinus ponderosa* | P05155 | 16 | seeds/$m^2$ | *Tilia cordata* | P05164 | 20 | seeds/$m^2$ |
| *Nothofagus pumilio* | D00002 | 18 | seeds/$m^2$ | *Nothofagus solandri* | P05004 | 8 | seeds/$m^2$ |
| *Quercus coccinea* | P05035 | 90 | seeds/$m^2$ | | | | |
| *Quercus spp.* | P05115 | 50 | g/$m^2$ | *Fagus crenata* | M04032 | 10 | g/$m^2$ |
| *Quercus ilex* | P05107 | 17 | g/$m^2$ | *Fagus sylvatica* | M00282 | 10 | g/$m^2$ |
| *Fagus sylvatica* | A00056 | 3 | g/$m^2$ | *Fagus sylvatica* | D06208 | 10 | g/$m^2$ |
| *Fagus sylvatica* | D06207 | 12 | g/$m^2$ | *Pinus sylvestris* | M01208 | 8 | g/$m^2$ |
| *Quercus spp.* | D00019 | 10 | seeds/ind. | *Quercus alba* | D00009 | 17 | seeds/ind. |
| *Quercus rubra* | D00009 | 16 | seeds/ind. | *Quercus montana* | D00009 | 17 | seeds/ind. |
| *Picea abies* | D00019 | 10 | seeds/ind. | *Abies alba* | D00019 | 10 | seeds/ind. |
| *Fagus sylvatica* | D00012 | 7 | seeds/ind. | *Pinus pinaster* | D00019 | 10 | seeds/ind. |
| *Quercus mongolica* | P05077 | 20 | seeds/ind. | *Pinus nigra* | D00019 | 10 | seeds/ind. |
| *Quercus petraea* | D00019 | 10 | seeds/ind. | *Quercus chapmanii* | P05001 | 60 | seeds/ind. |
| *Quercus geminata* | P05001 | 60 | seeds/ind. | *Quercus laevis* | P05001 | 30 | seeds/ind. |
| *Quercus myrtifolia* | P05001 | 60 | seeds/ind. | *Quercus wislizeni* | P05092 | 13 | seeds/ind. |
| *Quercus agrifolia* | P05092 | 63 | seeds/ind. | | | | |
| *Quercus ilex* | P05043 | 15 | seeds/branch | *Quercus pubescens* | P05043 | 15 | seeds/branch |
| *Quercus ilex* | M10139 | 50 | index | *Fagus grandifolia* | M01009 | 12 | index |
| *Acer saccharum* | M01009 | 12 | index | | | | |
| *Picea abies* | A00022 | 40 | cones/ind. | *Picea abies* | A00169 | 13 | cones/ind. |
| *Picea engelmannii* | P05047 | 20 | cones/ind. | *Pseudotsuga menziesii* | M01002 | 6 | cones/ind. |
| *Picea engelmannii* | D00024 | 56 | cones/ind. | *Pseudotsuga menziesii* | D00024 | 17 | cones/ind. |
| *Abies amabilis* | P05047 | 20 | cones/ind. | *Tsuga mertensiana* | P05047 | 20 | cones/ind. |
| *Pinus ponderosa* | D00038 | 5 | cones/ind. | *Picea glauca* | A00203 | 30 | cones/ind. |
| *Picea glauca* | P05096 | 100 | cones/ind. | *Abies lasiocarpa* | P05047 | 20 | cones/ind. |
| *Abies lasiocarpa* | D00024 | 33 | cones/ind. | *Abies grandis* | P05047 | 20 | cones/ind. |
| *Pinus albicaulis* | P05171 | 25 | cones/$m^2$ | *Abies lasiocarpa* | P05177 | 30 | cones/$m^2$ |
| *Pinus albicaulis* | D00041 | 3 | cones/branch | | | | |
| *Chionochloa rubra* | P05088 | 100 | flowers/ind. | | | | |
| *Quercus velutina* | M01012 | 18 | fruits/$m^2$ | *Quercus michauxii* | M01012 | 18 | fruits/$m^2$ |
| *Vaccinium vitis-idaea* | D00028 | 10 | fruits/$m^2$ | | | | |

### References


Bogdziewicz, M., Żywiec, M., Espelta, J. M., et al. (2019). Environmental veto synchronizes mast seeding in four contrasting tree species. *The American Naturalist*, 194(2):246–259.

Crone, E. E., McIntire, E. J. B., and Brodie, J. (2011). What defines mast seeding? Spatio-temporal patterns of cone production by whitebark pine. *Journal of Ecology*, 99(2):438–444.

LaMontagne, J. M., Greene, D. F., Holland, E. P., Johnstone, J. F., Schulze, M., et al. (2024). Community synchrony in seed production is associated with trait similarity and climate across North America. *Ecology Letters*, 27(12):e14498.

Loreau, M. and De Mazancourt, C. (2008). Species synchrony and its drivers: Neutral and nonneutral community dynamics in fluctuating environments. *The American Naturalist*, 172(2):E48–E66.

Satake, A. and Iwasa, Y. (2002). Spatially limited pollen exchange and a long-range synchronization of trees. *Ecology*, 83(4):993–1005.

Woodward, W. A., Sadler, B. P., and Robertson, S. (2022). *Time Series for Data Science: Analysis and Forecasting*. CRC Press, Boca Raton, FL.
